# A novel human blood-brain barrier model reveals pericytes as critical regulators of viral neuroinvasion

**DOI:** 10.1101/2024.12.18.629173

**Authors:** Alexsia Richards, Andrew Khalil, Punam Bisht, Troy W. Whitfield, Xinlei Gao, Roger Kamm, David Mooney, Lee Gehrke, Rudolf Jaenisch

**Affiliations:** Whitehead Institute for Biomedical Research, Cambridge, MA 02142; Wyss Institute for Biologically Inspired Engineering, Harvard University, Boston, MA 02115; John A. Paulson School of Engineering and Applied Sciences, Harvard University, Cambridge, MA, 02138; Department of Biological Engineering, Massachusetts Institute of Technology, Cambridge, MA, 02139; Department of Mechanical Engineering, Massachusetts Institute of Technology, Cambridge, MA, 02139; Department of Microbiology, Harvard Medical School, Boston, MA 02115; Institute for Medical Engineering and Science, Massachusetts Institute of Technology, Cambridge, MA, 02139; Department of Biology, Massachusetts Institute of Technology, Cambridge, MA, 02139

## Abstract

The blood-brain barrier (BBB) plays a vital role in regulating the passage of biomolecules between the bloodstream and the central nervous system (CNS) while also protecting the CNS from pathogens. Pericytes reside at the interface between the endothelial cells that form the vessel walls and the brain parenchyma. These cells are critical for maintaining BBB integrity and play key roles in regulating vessel permeability, blood flow, and immune cell migration. In this study, we developed a novel serum-free protocol to generate neural crest cell-derived pericytes (NCC-PCs) from human pluripotent stem cells (hPSCs). These NCC-PCs enhance BMEC barrier function and can be co-cultured with hPSC-derived brain microvascular endothelial cells (BMECs) in a contact co-culture BBB model that recapitulates the *in vivo* cellular interactions at the BBB. We used this model to evaluate the pathological consequences of BBB cell infection by highly neuroinvasive flaviviruses. Our results identify a previously undescribed role for NCC-PCs in maintaining BMEC barrier integrity during infection and reducing the spread of viral infection to the CNS.

## Introduction

Neuroinvasive flaviviruses, including West Nile virus (WNV), Powassan virus (POWV), and Japanese encephalitis virus (JEV), pose significant public health threats due to their ability to infect the central nervous system, leading to severe neurological complications^1^. The blood-brain barrier (BBB) is thought to be a key access point for these viruses to reach the CNS. Evidence from *in vitro* models has generated multiple hypotheses on how these viruses cross the BBB, including transcytosis through infected endothelial cells, disruption of the barrier, and transport via infection of migrating immune cells^2, 3^.

The BBB is a critical interface that tightly regulates the passage of biomolecules between the bloodstream and the central nervous system (CNS), while also protecting the CNS from invading pathogens. Pericytes (PCs) are multifunctional mural cells that wrap around the endothelial cells of capillaries and venules throughout the body^4^. At the BBB, the interaction between PCs and brain microvascular endothelial cells (BMECs) is crucial for regulating vessel permeability^5, 6^. Additionally, PCs can exhibit macrophage-like phagocytic phenotypes, contributing to the clearance of circulating toxins, and regulate the migration of immune cells across the BBB^7–9^.

Studying the mechanisms by which neuroinvasive viruses cross the BBB is difficult *in vivo* and has long been done with *in vitro* models. However, many *in vitro* models using primary cells often fail to incorporate multiple BBB cell types and exhibit poor endothelial barrier function, making it difficult to extrapolate these findings to *in vivo* pathology. Advancements in stem cell technology have improved *in vitro* BBB models by enabling the production of isogenic populations of nearly all BBB cell types^10^. During development, forebrain pericytes arise from neural crest cells (NCCs) ^11, 12^. Although protocols have been published for the differentiation of neural crest-derived pericytes (NCC-PCs) from human pluripotent stem cells (hPSCs) ^13, 14^, these protocols utilize high concentrations of serum in the differentiation media. In addition to its intrinsic variability, serum is known to activate inflammatory signaling and reduce endothelial barrier function, which presents challenges when incorporating these protocols with additional hPSC-derived BBB cell types^15, 16^. In addition, serum is known to inhibit infection with a number of viruses, further complicating studies on viral infection in these cells^17, 18^ . To address these potential issues, we developed a protocol for generating hPSC-derived NCC-PCs using fully defined, serum-free media for both differentiation and maintenance. We further integrated these NCC-PCs with BMECs to create an *in vitro* contact co-culture BBB model.

Using our novel hPSC-derived BBB model, we demonstrate that neuroinvasive flaviviruses infect multiple BBB cell types, while exerting divergent effects on BMEC barrier function that correlate with *in vivo* disease severity. Additionally, we identify a previously undescribed protective role for NCC-PCs during viral infection at the BBB.

## Results

### Serum-free differentiation of neural crest derived pericytes from hPSCs

We established a two-step protocol to generate and maintain neural crest derived PCs derived from hPSCs **(Fig. 1A)**. Similar to previously published protocols, our approach begins with the differentiation of NCCs from hPSCs^14, 19^ using a commercial kit (Stem Cell Technologies) following the manufacturer’s protocol. Following differentiation, NCCs were cultured in serum free pericyte media (sfPCM) for ten days to generate NCC-PCs. We have previously shown that sfPCM supports the differentiation and maintenance of mesoderm derived pericytes^20^. To confirm that our protocol worked with multiple hPSC lines we performed differentiation with both H1 embryonic stem cells and iPS11 induced pluripotent stem cells. We examined the expression of key marker genes at each step of our protocol by bulk RNA sequencing **(Fig. 1B-C, Supplemental Fig. 1)**. As expected, hPSCs expressed high levels of POU5F1 and NANOG, whereas NCCs had the highest expression of NGFR and SOX9 **(Fig. 1B, Supplemental Fig. 1A)**. We included both primary human brain PCs as well as hPSC derived mesoderm PCs as controls for PC gene expression. We examined a panel of genes known to be highly expressed in pericytes, including PDGFRa, PDGFRb, CSPG2 and ANGPT1 **(Fig. 1C, Supplemental Fig. 1B)**. Our NCC-PCs expressed these marker genes at levels comparable to primary pericytes and hPSC-derived mesoderm PCs **(Fig. 1C, Supplemental Fig. 1B)**. To further characterize our hPSC-derived NCC-PCs, we performed principal component analysis (PCA) on counts data from RNA-seq experiments using hPSC-derived NCC-PCs, hPSC-derived mesoderm PCs, and primary brain PCs **(Supplemental Fig. 2)**. In the PCA plot, the first principal component (PC1) captures nearly 41% of the variance and the second principal component (PC2) captures 20%. The first principal component delineates the initial hPSC cells (i.e. H1 hPSCs or iPS11 iPSCs) from the differentiated cells.

**Figure 1:**
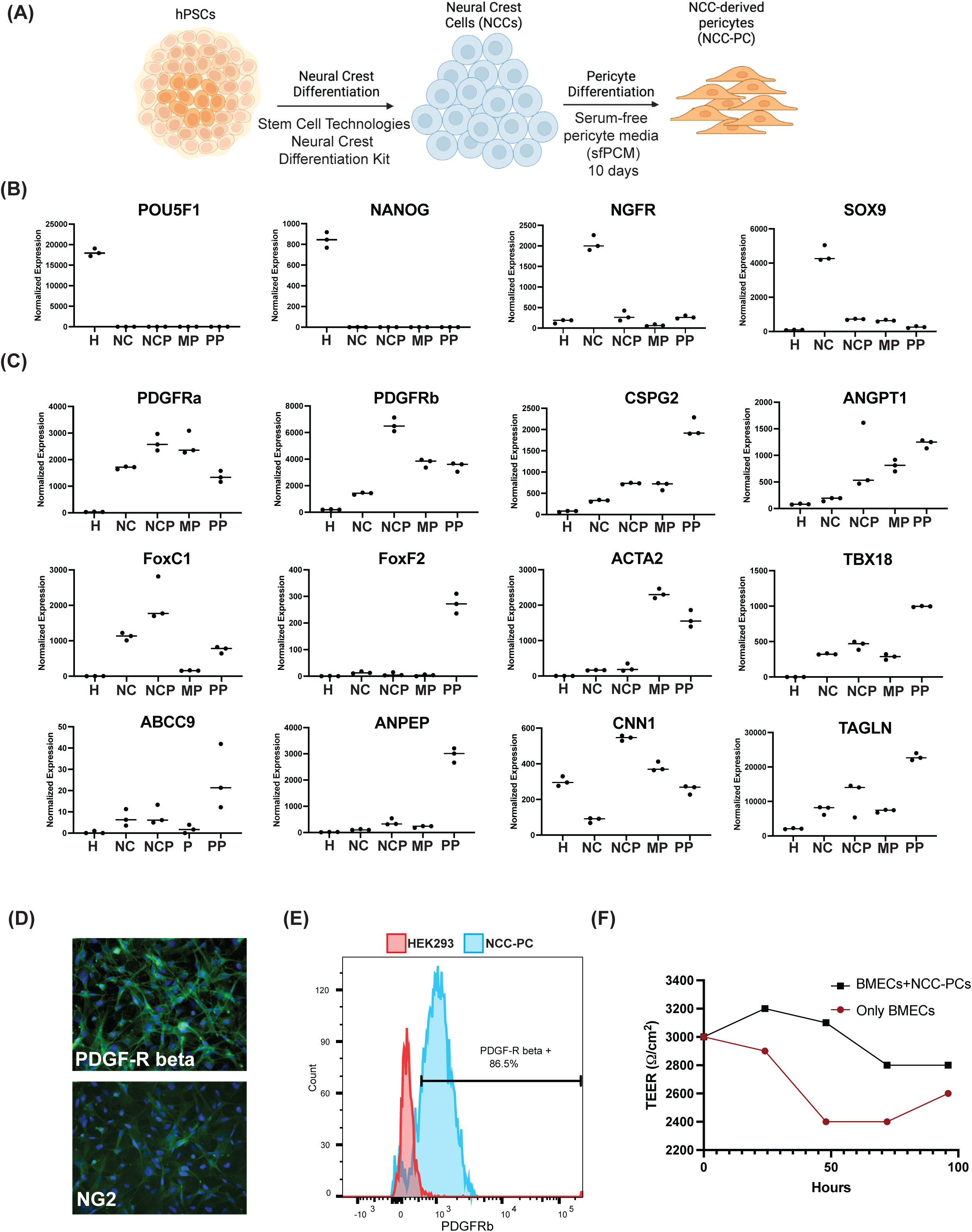
Generation and characterization of hPSC-derived neural crest pericytes. **(A)** Schematic of differentiation of neural crest pericytes (NCC-PCs) from hPSCs. **(B-C)** Bulk RNA sequencing was performed on hPSCs (H), hPSC-derived neural crest cells (NC), hPSC-derived NCC-PCs (NCP), hPSC-derived mesoderm pericytes (MP), or primary CNS pericytes (PP). H1 stem cells were used for all differentiated cells. **(B)** Normalized expression of hPSC marker genes POU5F1 and NANOG or neural crest marker genes NGFR and SOX9. **(C)** Normalized expression of pericyte marker genes. **(D)** Immunofluorescence images of NCC-PCs stained with antibodies directed again PDGF-R beta or NGD. **(E)** Expression of PDGF-R beta was quantitated by flow cytometry on NCC-PCs or control HEK293 cells. **(F)** TEER quantitation of BMECs plated in Transwell plates with or without NCC-PCs plated in the lower chamber. TEER was measured beginning at 0 hours after the initiation of co-culture.

Immunohistochemistry showed that NCC-PCs expressed PDGFRb and NG2 **(Fig. 1D)** and were greater than 85% positive for PDGFRb relative to control HEK293 cells (**Fig. 1E)**. The co-culture of brain microvascular endothelial cells (BMECs) with PCs is known to improve barrier function^19^. Transendothelial electrical resistance (TEER) provides a quantitative read out of endothelial cell barrier function^21^ . To determine if co-culture of NCC-PCs with BMECs could improve BMEC barrier function, hPSC-derived BMECs were plated in the apical chamber of a Transwell plate and NCC-PCs in the lower chamber. TEER values were measured every 24 hours for up to 96 hours after the initiation of co-culture, our results show that co-culture with NCC-PCs increased BMEC TEER values at all time points examined **(Fig. 1F)**. In conclusion, our data shows that functional NCC-PCs can be derived from hPSCs using fully-defined serum free conditions. The use of serum-free pericyte media for differentiation is particularly critical as we observed that even low concentrations of serum in the media result in a significant reduction in BMEC barrier integrity **(Supplemental Fig. 3)**.

### Neuroinvasive flaviviruses infect hPSC-derived blood brain barrier cells

To determine if hPSC-derived BBB cells can be productively infected by neuroinvasive flaviviruses hPSC-derived BMECs, astrocytes, and NCC-PCs were infected with WNV, POWV, or JEV at an MOI of 0.1 and the amount of infectious virus released into the media quantitated at 0 hours, 24 hours, or 48 hours post-infection. We observed robust replication of all three viruses in all three hPSC-derived cell types **(Fig. 2A)**. One possible mechanism of spread of infection through the BBB is via basolateral release from infected BMECs to the surrounding perivascular CNS cells. To determine if infected BMECs released virus basolaterally following apical infection, BMECs were cultured in Transwell plates and infected apically with WNV, JEV, or POWV. The amount of infectious virus released into the lower chamber was quantified at 24 and 48 hours **(Fig. 2B)**. Although infection with all three viruses resulted in basolateral release of nascent virus to the lower chamber, POWV infection resulted in significantly more infectious virus in the lower chamber at 24 hours relative to WNV or JEV **(Fig. 2B)**.

**Figure 2:**
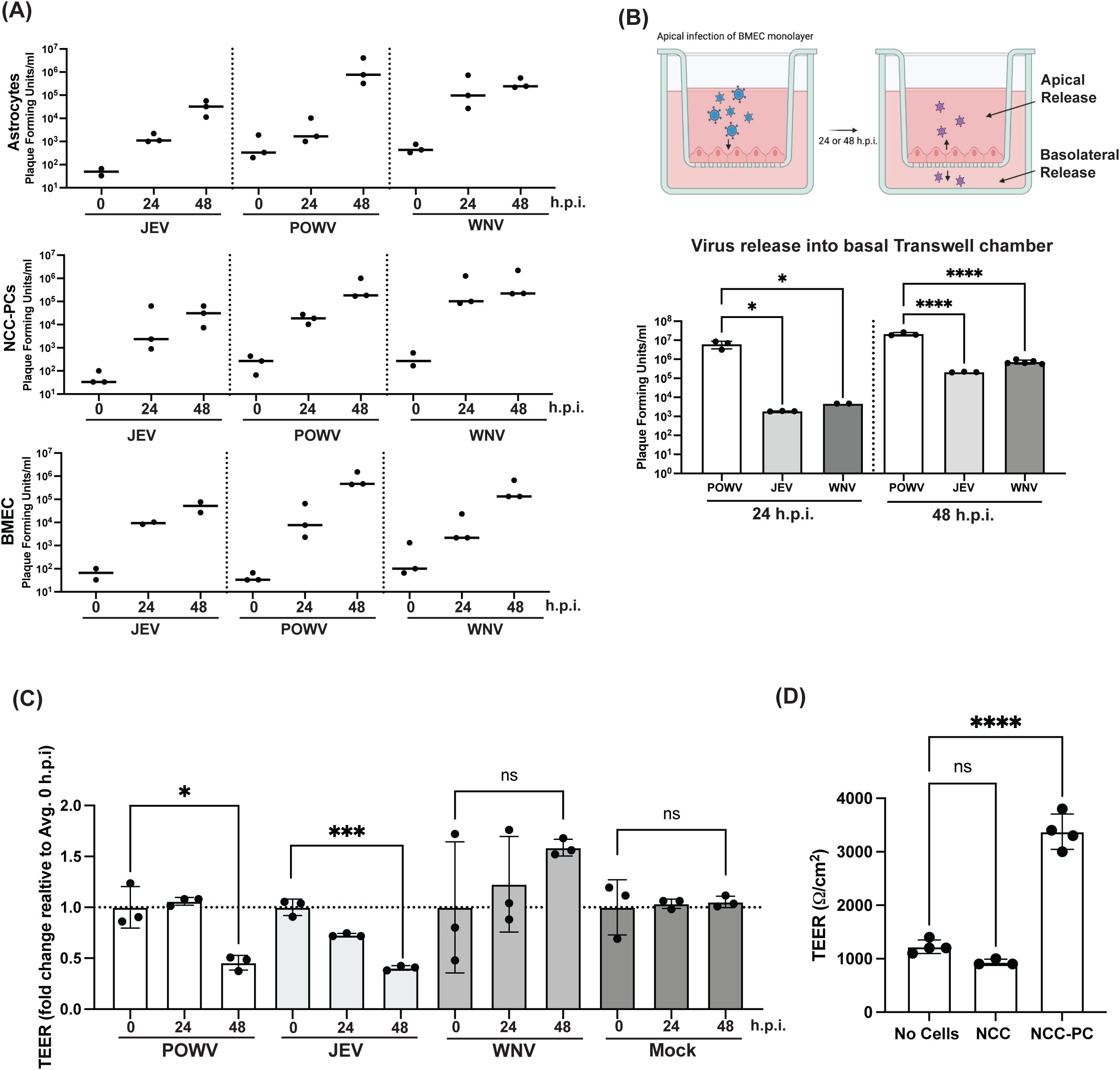
Neurotropic flaviviruses productively infect hPSC-derived blood brain barrier cells. **(A)** hPSC-derived astrocytes, NCC-PCs, or BMECs were infected with either JEV, POWV, or WNV at an MOI of 0.1 and the amount of infectious virus released into the media at the indicated time post-infection was quantitated by plaque assay. **(B)** hPSC-derived BMECs were plated in Transwell plates and apically infected with WNV, POWV, or JEV (MOI=0.1). The amount of infectious virus released into the basal chamber of the Transwell was quantitated by plaque assay. **(C)** hPSC-derived BMECs were plated in Transwell plates and TEER measured at the indicated time after infection with WNV, POWV, or JEV. **(D)** BMECs were plated Transwell plates with either hPSC-derived NCCs or NCC-PCs plated in the lower chamber. BMECs were infected with POWV 24 hours after the initiation of co-culture and TEER measured at 72 hours post-infection.

We next examined TEER values for BMECs cultured in Transwell plates at 24, 48, or 72 hours post infection. WNV and JEV infection did not reduce BMEC barrier integrity relative to mock infected BMECs **(Fig.2C)**. Conversely, beginning at 48 hours after infection with POWV we observed a significant reduction in BMEC TEER **(Fig. 2B)**. Our previous data showed that co-culture of NCC-PCs with BMECs improved barrier function **(Fig. 1F)**. To determine if co-culture with NCC-PCs could improve BMEC barrier function following POWV infection, BMECs were plated in the apical chamber of a Transwell plate and NCC-PCs or their precursor NCCs were plated in the lower chamber. BMECs were infected with POWV 24 hours after the initiation of co-culture and TEER measured at 72 hours post-infection. The co-culture of BMECs with NCC-PCs but not their precursor NCCs resulted in improved BMEC barrier function following POWV infection **(Fig. 2D)**.

In summary, our data show that the cell types that comprise the BBB can be infected with multiple neuroinvasive flaviviruses and infected BMECs release nascent virus both apically and basolaterally. Collectively, these data support the hypothesis that infection at the BBB is a viable route for these viruses to enter the CNS. In addition, our data show that POWV may be unique amongst flaviviruses in its rapid impact on BMEC barrier integrity. The increased basolateral release of POWV coupled with disruption of the BMEC barrier function may in part contribute to the increased fatality rate of POWV relative to WNV or JEV^22–24^.

### A contact co-culture BBB model reveals a role for pericytes in reducing WNV neuroinvasion

BBB function is highly dependent on the interactions between the endothelial cells that make up the vascular wall and the perivascular cells that surround them. The generation and co-culture of multiple isogenic populations of BBB cells is a key advantage of hPSC-derived BBB models. The Transwell culture system is comprised of two culture compartments separated by a semipermeable membrane **(Fig. 3A)**. Previous hPSC-derived BBB models have cultured BMECs on the semipermeable membrane and plated mural or brain cells in the lower chamber^14, 19^. This set up enables paracrine signaling between cell types, but may not recapitulate interactions that require the close proximity or contact between BMECs and pericytes or glial cells. To improve on current hPSC-derived Transwell BBB models we tested if mural and glial cells could be cultured on the basolateral side of the same Transwell membrane that BMECs were seeded **(Fig. 3A)**. Two days after BMEC seeding, the insert was inverted and Matrigel was added to the basolateral side of the membrane. Following Matrigel addition hPSC-derived NCC-PCs or hPSC-derived astrocytes were added to the basolateral side of the membrane. Cells were allowed to adhere for 3 hours, at which point the Transwell insert was placed back in a standard culture plate and media added to both compartments. Fluorescently-tagged NCC-PCs or astrocytes were used to enable easy confirmation of successful seeding of the basolateral side of the Transwell membrane by live cell imaging **(Fig. 3B)**. We refer to this novel contact co-culture BBB model as the ccBBB model.

**Figure 3:**
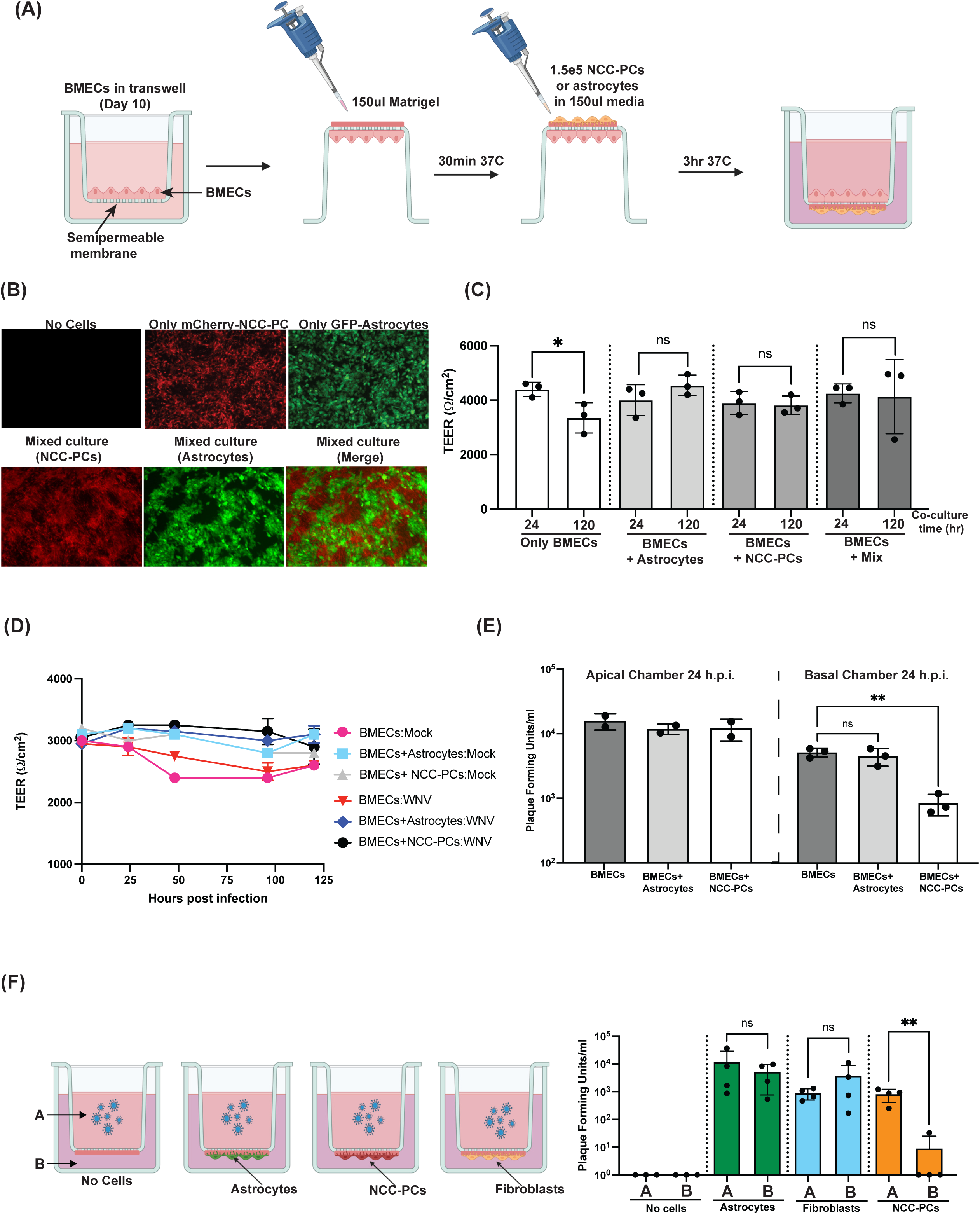
Generation and infection of a contact co-culture blood brain barrier model (ccBBB). **(A)** Schematic of generation of hPSC-derived contact co-culture blood brain barrier model (ccBBB). **(B)** Images of mCherry-tagged NCC-PCs or GFP-tagged astrocytes cultured on the underside of Transwell semipermeable membrane in ccBBB system. **(C)** TEER measurements across Transwell semipermeable membrane of ccBB system. **(D)** Transcytosis of FITC–tagged 10-kDa dextran across the ccBBB membrane. FITC-dextran was added to the upper chamber and the fluorescence intensity of media the lower chamber was quantitated four hours after FITC-dextran addition. All results are expressed as a fold change relative to the average value ccBBB system with no cells. **(E)** Schematic of ccBBB conditions and infection procedure. **(F)** TEER measurements across ccBBB membrane at the indicated time after infection with WNV or mock infection. **(G)** WNV infection of ccBBB system with only BMECs, or BMECs with NCC-PCs or astrocytes. WNV infection was performed 72 hours after the initiation of co-culture. The amount of infectious virus in the top and bottom chamber at 24 hours post infection was quantitated by plaque assay.

To identify the optimal co-culture conditions for our ccBBB model we examined the effect of different media compositions in the basolateral chamber. We observed that prolonged astrocyte co-culture was significantly improved by replacing the media in the lower chamber with astrocyte media rather than sfPCM **(Supplemental Fig. 4A)**. As our goal was to be able to culture both mural and glial cells on the basolateral side of the membrane using the same culture conditions, we confirmed that NCC-PCs retained expression of PC specific genes when cultured in astrocyte media. We examined expression of the PC markers PDGFRb and NG2 following seven days of culture in either astrocyte media or sfPCM by immunofluorescence. Our results showed similar expression patterns in both conditions, suggesting that the NCC-PCs retained their cellular identity when cultured in astrocyte media **(Supplemental Fig. 4B)**.

In agreement with previous reports showing that co-culture with pericytes or astrocytes can improve BMEC barrier function^19, 25^, the addition of either NCC-PCs or astrocytes to the basolateral side of the membrane resulted in increased TEER values across the BMEC monolayer **(Fig. 3C)**. Critically, in the absence of BMECs neither NCC-PCs or astrocytes showed TEER values above baseline suggesting that the observed increase in TEER in the co-culture conditions was the due to the influence of these cells on BMECs **(Fig. 3C, Supplemental Fig. 4C)**. We also examined the impact of NCC-PCs or astrocytes on ccBBB permeability to 10kDa FITC-dextran. FITC-dextran was dosed into the apical Transwell chamber and the amount of dextran reaching the lower chamber was quantitated relative to the BMEC only condition **(Supplemental Fig. 4C).** The addition of astrocytes or fibroblasts to the ccBBB system did not impact BMEC permeability to dextran relative to the BMEC only condition, whereas the addition of NCC-PCs resulted in an 85% reduction in dextran permeability relative to the BMEC only condition **(Fig. 3D)**. These data suggest that NCC-PCs may have a more pronounced impact on reducing the leakiness of BMECs at the BBB than do astrocytes or fibroblasts.

In our ccBBB system, the apical side of the Transwell represents the blood, while the basolateral chamber represents the CNS. To determine if pericytes or astrocytes influence barrier integrity of release of virus at the BBB during viral infection, WNV was added to the apical side of the Transwell to initiate infection of the BMEC monolayer **(Fig. 3E)**. In agreement with the data presented in Fig. 2B we did not observe a significant reduction in TEER due to WNV infection relative to mock infection in any of the conditions tested **(Fig. 3F)**. The addition of NCC-PCs or astrocytes did not impact the apical release of virus from the BMEC monolayer, however, the addition of NCC-PCs to the system resulted in nearly a 10-fold decrease in the amount of virus released to the basolateral chamber relative to the BMEC only control at 24 hours post infection **(Fig. 3G)**. The addition of astrocytes had no effect on the amount of virus present in the basolateral chamber suggesting that the ability to dampen basolateral viral release may be a specific function of pericytes at the BBB.

### BMECs and pericytes display divergent early transcriptional responses to WNV BBB infection

When circulating virus reaches the BBB it is likely that the first cells exposed to virus are the BMECs that make up the vessel walls, followed by the mural or glial cells that surround these vessels. There is extensive cross-talk between cells at the BBB which has the potential to influence an individual cell’s response to infection or influence cells that are in close proximity to infected cells. For this reason, analyzing the transcriptional profile of BBB cells in isolation likely does not provide an accurate picture of the response to infection. To gain insight into the transcriptional response of BMECs and PCs at the BBB at discreet time points after exposure to WNV, we performed bulk RNA sequencing on BMECs and NCC-PCs cultured in the ccBBB system following apical infection **(Fig. 4A)**. We examined the response at 8 hours post infection, a time point where we speculated that infection would not have spread from the BMEC monolayer to NCC-PCs. We also examined the response at 48 hours post infection, a time point where we hypothesized that both cell populations would be infected. Critically, the gene expression patterns at each time point were compared to mock infected BMECs and NCC-PCs cultured in the ccBBB system, therefore the observed changes in gene expression were likely not due to culture conditions but were instead a specific result of viral infection.

**Figure 4:**
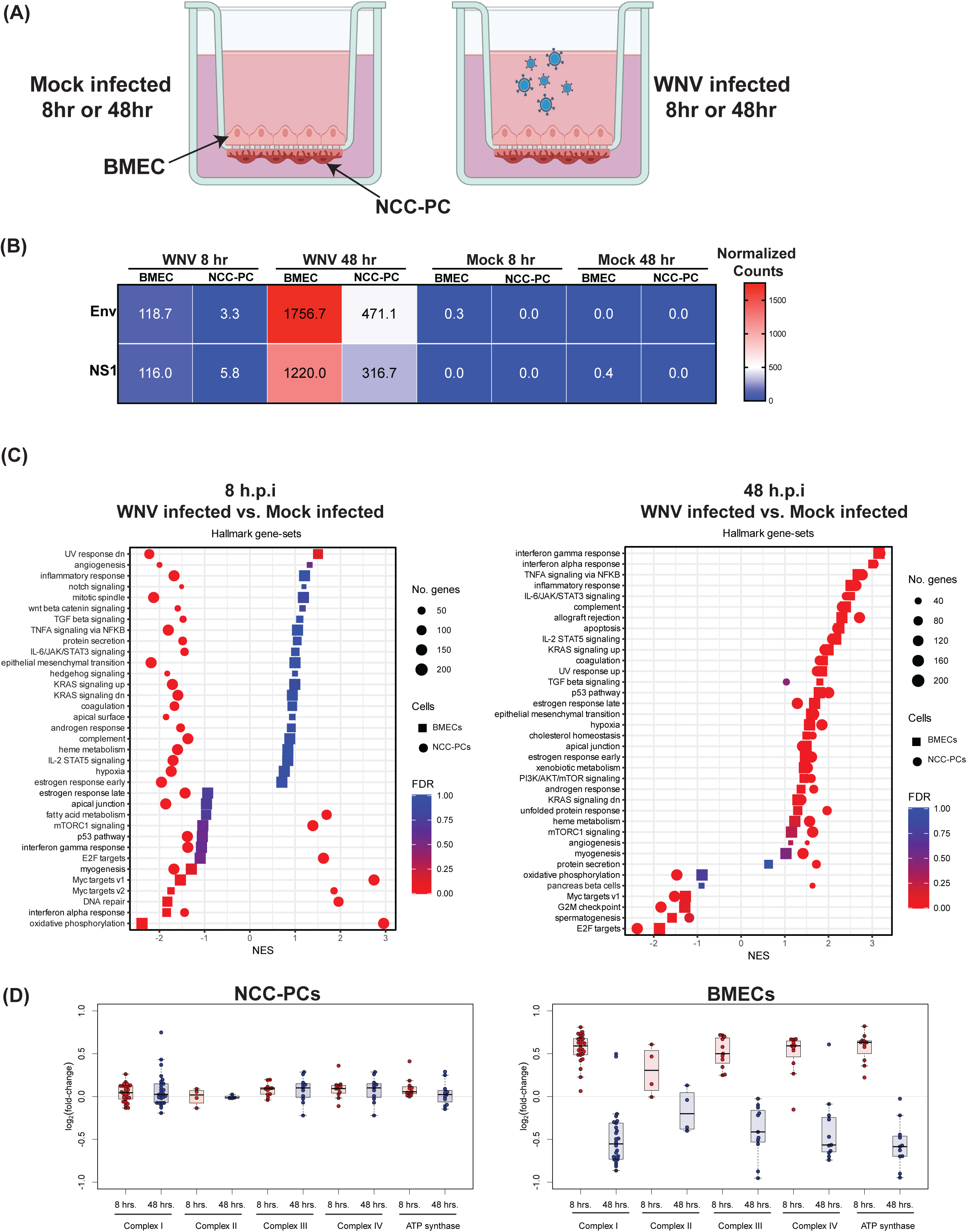
Transcriptional response to infection of BMECs and NCC-PCs cultured in the ccBBB system. **(A)** hPSC derived BMECs and NCC-PCs were plated in the ccBBB model as described in Fig. 3A. WNV was added to the apical chamber at an MOI of 1. Total cellular RNA was isolated from BMECs and NCC-PCs at the indicated time post infection and analyzed by bulk RNA sequencing. **(B)** RNA-seq counts, normalized via the median-of-ratios method in DESeq2^32^, for the Env and NS1 regions of the WNV reference genome following mock-or WNV-infection of the ccBBB model. **(C)** Gene set enrichment analysis (GSEA)^62^ was performed to compare WNV-infected versus mock-infected BMECs and WNV-infected versus mock-infected NCC-PCs at 8 and 48 hours post infection, respectively. The Hallmark collection^32^ of gene-sets from the MSigDB was used and gene-sets were plotted only when FDR < 0.05 for enrichment in at least one of the between-condition comparisons. **(D)** Average differential gene expression for genes the encode for proteins in the four complexes of the electron transport chain and ATP synthase. In the ccBBB model, log_2_(fold-change) was first estimated from RNA-seq data using DESeq2^32^ by comparing WNV infected versus mock infected NCC-PCs and WNV infected versus mock infected BMECs, respectively, with statistical comparisons subsequently made over all genes in the indicated metabolic pathways (see Table 1) using one-sample t testing.

At 8 hours after virus exposure the NCC-PCs exhibited very low levels of WNV transcription: the fraction of normalized reads mapping to either the Env or NS1 regions of the WNV genome was below 2% of its level at 48 hours post-exposure and just 5% of the level for BMECs at 8 hours post-exposure **(Fig. 4B)**. Our data suggest that the BMEC monolayer remains intact during infection **(Fig. 3F)**, which led to the hypothesis that any virus that reached the NCC-PCs had to go through the BMEC monolayer to reach the NCC-PCs **(Fig 4A)**. Thus, the viral RNA detected in NCC-PCs could be the result of early release of progeny virions from infected BMECs or transcytosis of input virus in between the tight junctions of the BMEC monolayer.

In BMECs at 8 hours post infection, the overall changes in gene expression upon infection were small with few individual genes significantly different from mock infected BMECs **(Supplemental Fig. 5A-B)**. Through gene set enrichment analysis (GSEA)^26^, we observed a downregulation in the IFN-α and oxidative phosphorylation gene-sets **(Fig. 4C)**. By 48 hours post infection both BMECs and NCC-PCs showed strong upregulation of inflammatory signaling pathways, including IFN-α and IFN-γ responses and TNFα signaling **(Fig. 4C Supplemental Fig. 5C-D)**. These results are consistent with a robust innate immune response to infection and suggest that during infection at the BBB both BMECs and pericytes contribute to the local innate immune response.

At 8 hours post infection NCC-PCs showed significant downregulation of a number of genes with a notable exception of HOXA1 which was significantly upregulated (**Supplemental Fig. 5B)**. GSEA analysis on NCC-PCs 8 hours after WNV exposure showed that the majority of gene-sets including those involved in IFN-α and IFN-γ responses were downregulated **(Fig. 4C)**. Surprisingly, the oxidative phosphorylation gene-set was strongly upregulated in NCC-PCs at 8 hours post infection. This result was particularly notable as the oxidative phosphorylation gene-set was one of the most downregulated gene sets in the BMECs at 8 hours after virus exposure. To further explore the specific genes driving upregulation of this gene set we examined the differential expression of genes that produce the four large complexes that comprise the electron transport chain as well as ATP synthase^27^. We examined expression of these genes in NCC-PCs at both 8 and 48 hours after virus exposure to determine how expression levels changed over the course of infection. Our results show that the expression of genes encoding for Complexes I through Complex IV and ATP synthase is upregulated in NCC-PCs at 8 hours post infection relative to mock infected NCC-PC (µ_log (fold_ = 0.53, p-value = 3.7 x 10^-30^), whereas by 48 hours post infection the average expression of these genes is down in infected NCC-PCs relative to mock infected NCC-PCs (µ_log (fold_ = -0.46, p-value = 4.3 x 10^-is^, see **Fig. 4D**, **Table 1**). Conversely, WNV infection of BMECs resulted in much smaller changes in the expression level of these genes relative to mock infection **(Fig. 4D**, **Table 1)**, suggesting that the shift in expression of metabolic genes may be unique feature of NCC-PCs.

**Table 1.**

|  | NCC-PC<br>8 h.p.i. | NCC-PC<br>48 h.p.i. | EC<br>8 h.p.i. | EC<br>48 h.p.i. |
| --- | --- | --- | --- | --- |
| **COMPLEX**<br>**I** | **Log2FC (WNV infected vs. Mock infected)** |  |  |  |
| NDUFA3 | 0.73415583 | -0.735119 | 0.12301695 | 0.43124366 |
| NDUFS7 | 0.72121634 | -0.7320411 | 0.05357259 | 0.0148358 |
| NDUFA5 | 0.54199786 | -0.2290166 | -0.1336208 | -0.0736978 |
| NDUFV1 | 0.54765869 | -0.4900453 | 0.16868784 | -0.0190035 |
| NDUFS8 | 0.66625103 | -0.5322549 | 0.11984612 | 0.04564935 |
| NDUFB1 | 0.81028164 | -0.6963036 | -0.0055606 | 0.23925238 |
| NDUFS6 | 0.75361237 | -0.7092134 | 0.11976456 | -0.0074054 |
| NDUFS2 | 0.52376423 | -0.3101402 | -0.0143121 | -0.1924391 |
| NDUFB7 | 0.70723726 | -0.6308093 | 0.26131404 | 0.19612287 |
| NDUFB5 | 0.43610581 | -0.2854488 | -0.1002111 | 0.07446164 |
| NDUFC1 | 0.62065097 | -0.8137267 | -0.029226 | 0.173205 |
| NDUFB4 | 0.62828558 | -0.783272 | 0.05694262 | 0.00689658 |
| NDUFV2 | 0.61541306 | -1.5172208 | 0.12497301 | 0.36997966 |
| NDUFA2 | 0.56648721 | -0.6566156 | 0.08063125 | -0.0740509 |
| NDUFB2 | 0.73549161 | -0.5732605 | 0.14714191 | 0.10586018 |
| NDUFAB1 | 0.63370392 | -0.3622364 | 0.14633021 | 0.02894066 |
| NDUFA8 | 0.42065541 | -0.4007474 | 0.1307616 | -0.1004056 |
| NDUFB6 | 0.675851 | -0.7525585 | -0.0845667 | 0.02123415 |
| NDUFB3 | 0.67129141 | -0.7048174 | -0.0594731 | 0.1494488 |
| NDUFA1 | 0.4980123 | -0.4961723 | 0.08555226 | 0.14061784 |
| NDUFC2 | 0.48693871 | -0.4270126 | -0.1318622 | -0.009632 |
| NDUFS3 | 0.32206844 | -0.2984467 | 0.03398795 | -0.0754295 |
| NDUFA9 | 0.52576355 | 0.49739389 | 0.03638763 | 0.7485592 |
| NDUFB8 | 0.48576734 | -0.8646578 | -0.0097366 | -0.0729838 |
| NDUFA7 | 0.22700329 | -0.2027933 | 0.02679368 | 0.07615677 |
| NDUFS1 | 0.06373369 | 0.46655682 | -0.0638568 | -0.1289948 |
| **COMPLEX**<br>**II** |  |  |  |  |
| SDHD | 0.46727013 | -0.0399384 | -0.1352324 | -0.018834 |
| SDHB | 0.60830476 | -0.3599651 | 0.08891354 | 0.02407335 |
| SDHA | 0.14562393 | -0.4001886 | 0.05337868 | -0.0127022 |
| SDHC | -0.005411 | 0.1300804 | -0.0164706 | -0.0198931 |
| COMPLEX III |  |  |  |  |
| UQCR10 | 0.67070124 | -0.5548368 | -0.04022 | 0.12963891 |
| UQCRC1 | 0.50050766 | -0.5088408 | 0.1962969 | 0.13881084 |
| UQCRB | 0.71491723 | -0.8746007 | 0.02915534 | 0.16153169 |
| UQCRQ | 0.69808171 | -0.4813526 | 0.09859697 | 0.26074117 |
| UQCRH | 0.71845915 | -0.9529868 | 0.19275587 | -0.0257976 |
| UQCRFS1 | 0.47317318 | -0.2185043 | -0.0125806 | 0.10068459 |
| UQCRC2 | 0.34529886 | -0.0252204 | 0.09490501 | 0.28639341 |
| UQCR11 | 0.43217798 | -0.413581 | 0.02507798 | -0.07088 |
| CYB5A | 0.24890995 | -0.1944255 | 0.07461507 | 0.07216987 |
| CYB5R3 | 0.25683909 | -0.0987683 | 0.11357212 | -0.2211945 |
| CYCS | 0.57952169 | -0.1280589 | 0.09055662 | 0.00647981 |
| COMPLEX IV |  |  |  |  |
| COX8A | 0.66761701 | -0.4698427 | 0.12606567 | 0.12963891 |
| COX6C | 0.6658216 | -0.7407538 | 0.13326632 | 0.13881084 |
| COX6B1 | 0.67176742 | -0.5649744 | 0.09266763 | 0.16153169 |
| COX5B | 0.53852761 | -0.2214017 | 0.14243971 | 0.26074117 |
| COX5A | 0.59532008 | -0.5725552 | -0.0174467 | -0.0257976 |
| COX6A1 | 0.59171239 | -0.6355804 | 0.09178922 | 0.10068459 |
| COX17 | 0.63230174 | -0.6566795 | 0.35969916 | 0.28639341 |
| COX7A2L | 0.38849774 | -0.088079 | 0.05144217 | -0.07088 |
| COX7C | 0.57034503 | -0.7001174 | 0.02693581 | 0.07216987 |
| COX11 | 0.26717671 | -0.2680604 | -0.1120233 | -0.2211945 |
| COX10 | -0.1521718 | 0.60814815 | 0.10663175 | 0.00647981 |
| ATP Synthase |  |  |  |  |
| ATP5MC2 | 0.57848386 | -0.4800649 | 0.04341081 | 0.02250038 |
| ATP5F1C | 0.65546633 | -0.6963364 | 0.09376902 | -0.0025823 |
| ATP5MC1 | 0.82062426 | -0.9011945 | 0.05557779 | -0.1123253 |
| ATP5PD | 0.61037096 | -0.5723158 | 0.10155453 | 0.05430539 |
| ATP5MC3 | 0.63220544 | -0.4447375 | 0.01224289 | -0.0308854 |
| ATP5F1E | 0.64426379 | -0.6276007 | 0.12724126 | 0.28964142 |
| ATP5PF | 0.63426113 | -0.6986329 | 0.02277773 | 0.04527836 |
| ATP5MF | 0.70758693 | -0.5845765 | 0.18514455 | 0.24961349 |
| ATP5F1B | 0.35948702 | -0.2202548 | 0.04488177 | -0.146352 |
| ATP5F1A | 0.22225761 | -0.0254319 | -0.0003634 | -0.0951949 |
| ATP5F1D | 0.42308395 | -0.9463665 | 0.41109238 | 0.08927844 |

## Discussion

The BBB is perhaps one of the most effective innate defense mechanisms our body has to protect the brain, which must maintain a delicate balance between maintaining the health of neural cells, mitigating the damage induced by excessive inflammation, and fighting a viral infection. The devastating consequences of damage to the BBB were highlighted by recent data showing that in patients suffering from long COVID disruption of the BBB correlates with cognitive impairment ^28^. Our data presented here shows that pericytes, which are known targets for SARS-CoV-2^20^ infection can function as critical regulations of BBB disruption during infection at the BBB.

In this study, we developed a novel, serum-free protocol to produce neural crest-derived pericytes (NCC-PCs) from hPSCs and incorporated them into a blood-brain barrier (BBB) model. Our model successfully mimics *in vivo* BBB anatomy, where brain microvascular endothelial cells (BMECs) and pericytes share a basement membrane, providing a more physiologically relevant platform for studying BBB interactions and viral neuroinvasion.

We used this novel BBB model to investigate how pericytes contribute to viral control within the BBB. Specifically, our findings suggest that pericytes may dampen the spread of WNV from BMECs to the brain. Our data showed that NCC-PCs are susceptible to viral infection, thereby raising the important question of how these cells could be reducing the release of virus to the lower chamber. Given that apical release of nascent virus from BMECs is not impacted (Fig. 3F), it is unlikely that these cells significantly reduce BMEC infection. However, it is possible that NCC-PCs specifically dampen basolateral release from BMECs. Alternatively, NCC-PCs may have reduced basolateral release of the virus compared to BMECs or astrocytes and therefore once infected are inefficient at spreading the infection to subsequent cells in the CNS.

This study is the first to examine the transcriptional response of the cells that comprise the BBB in a physiological setting where intact paracrine signaling occurs between the cell types during infection. By maintaining the natural interactions between BMECs, pericytes, and astrocytes, our model provides a more accurate representation of the *in vivo* environment, allowing for a deeper understanding of how these cells collectively respond to viral invasion. We observed that the transcription factor HOXA1 was significantly upregulated in pericytes early after WNV infection of neighboring BMECs. HOXA1 has been linked to the trans-differentiation of vascular smooth muscle cells into macrophage-like cells^29^. Pericytes have also been reported to adopt macrophage-like properties ^30^. It is possible that upregulation of HOXA1 may promote this transition when the pericytes as exposed to nearby infected cells. This response, alongside metabolic changes, could represent a critical mechanism by which pericytes protect the brain during viral infection. While our studies focused primarily on the role of neural crest derived pericytes at the BBB, it is possible that other mural cell types also play key roles in dampening the spread of infection at the BBB. Future studies incorporating smooth muscle cells and mesoderm derived pericytes into the ccBBB system will test the cell-type specific effects of brain mural cells during viral infection at the BBB.

We also observed that early after infection of BMECs nearby NCC-PCs increase expression of genes involved in oxidative respiration. It has been proposed that pericytes switch to oxidative metabolism to promote terminal differentiation and restrict proliferation^31^. Future studies are needed to investigate if during viral infection at the BBB changes in NCC-PC oxidative phosphorylation are protective or pathogenetic. Overall, our findings suggest that pericytes play an essential role in modulating viral infection at the BBB, providing new insights into how these cells can be leveraged for therapeutic interventions targeting viral neuroinvasion.

## Acknowledgments

We acknowledge funding from the Wyss Institute, and the Whitehead Institute. We thank the Genome Technology Core and Keck Imaging Facility at Whitehead Institute for Biomedical Research. This work was supported by NIH grant U19AI131135, NIBIB grant T32 EB016652. This work was performed in part in the Ragon Institute BSL3 core, which is supported by the NIH-funded Harvard University Center for AIDS Research (P30 AI060354) and the Massachusetts Consortium on Pathogen Readiness (MassCPR).

---

**Figure S1:**
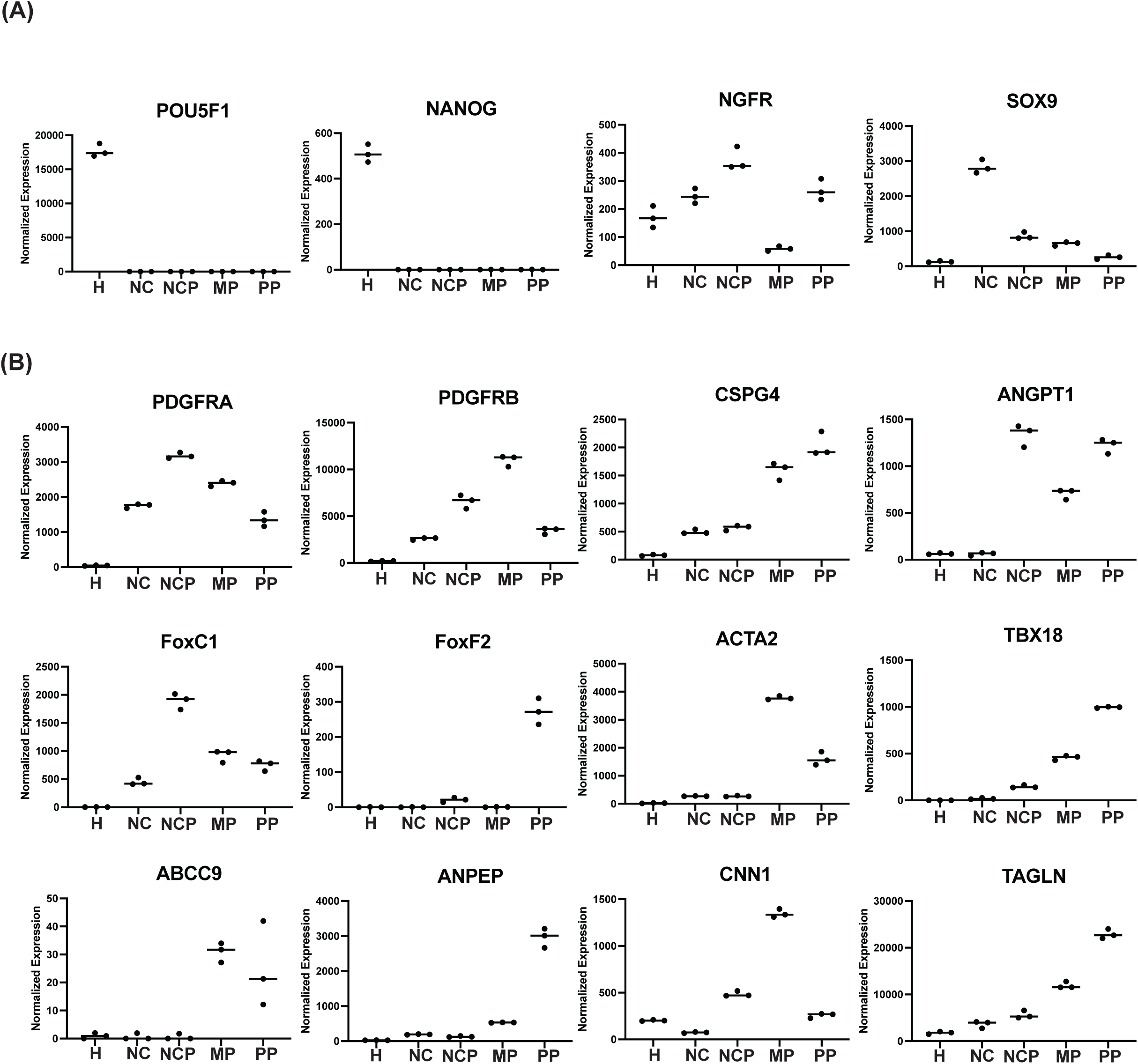
Characterization of hPSC-derived neural crest pericytes. Bulk RNA sequencing was performed on hPSCs (H), hPSC-derived neural crest cells (NC), hPSC-derived NCC-PCs (NCP), hPSC derived mesoderm pericytes (MP), or primary CNS pericytes (PP). iPS-11 stem cells were used for all differentiations. **(A)** Normalized expression of hPSC marker genes POU5F1 and NANOG or neural crest marker genes NGFR and SOX9. **(B)** Normalized expression of pericyte marker genes.

**Figure S2:**
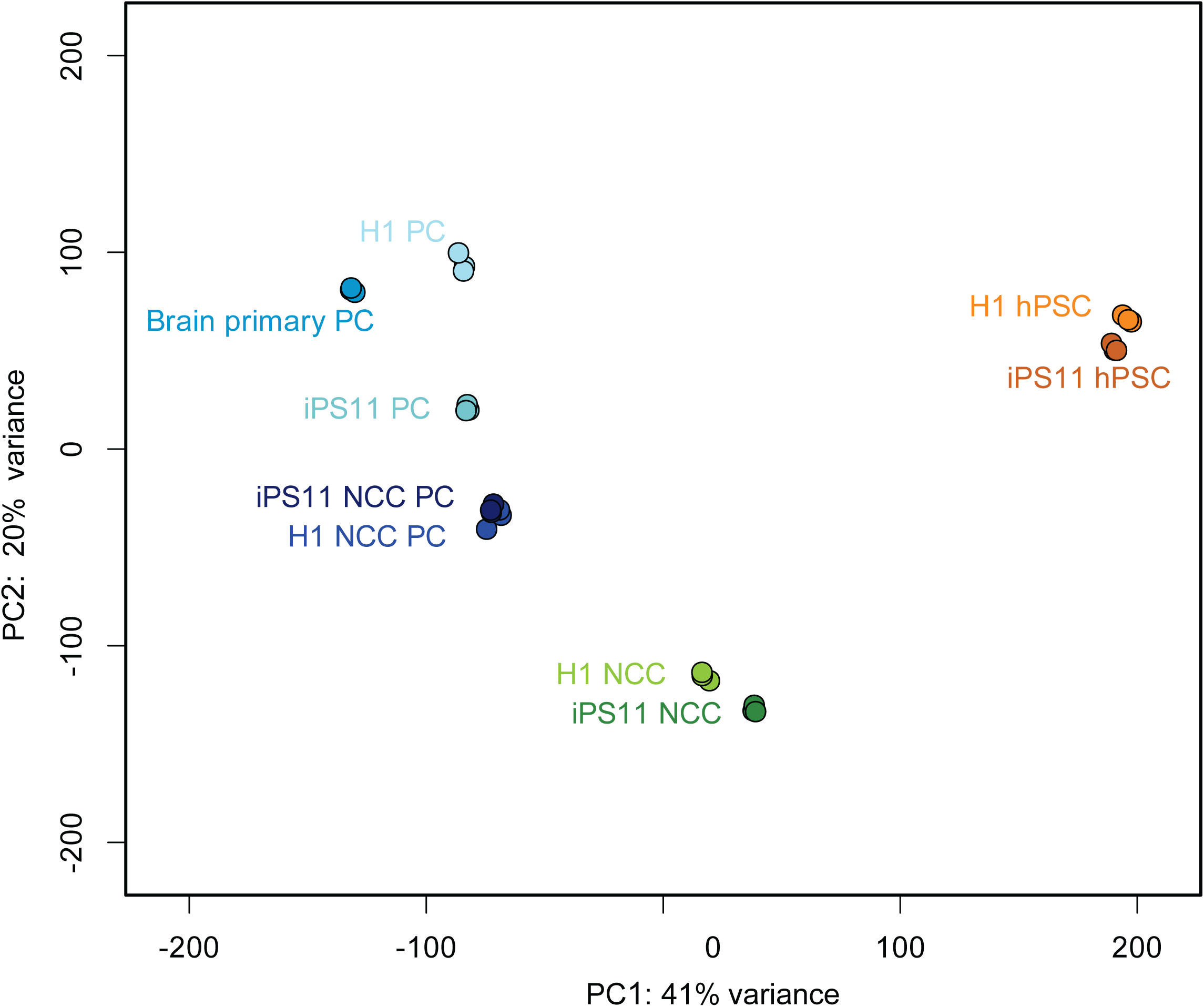
Principal component analysis of hPSC-derived pericytes and primary neural pericytes. Principal component analysis (PCA) on bulk-RNA sequencing data from hPSCs, hPSC-derived neural crest cells (NCC), hPSC-derived neural crest pericytes (NCC-PC), hPSC-derived mesoderm pericytes (PC), and primary brain vascular pericytes. Either H1 or iPS11 hPSCs were used for all differentiations as indicated in the figure.

**Figure S3:**
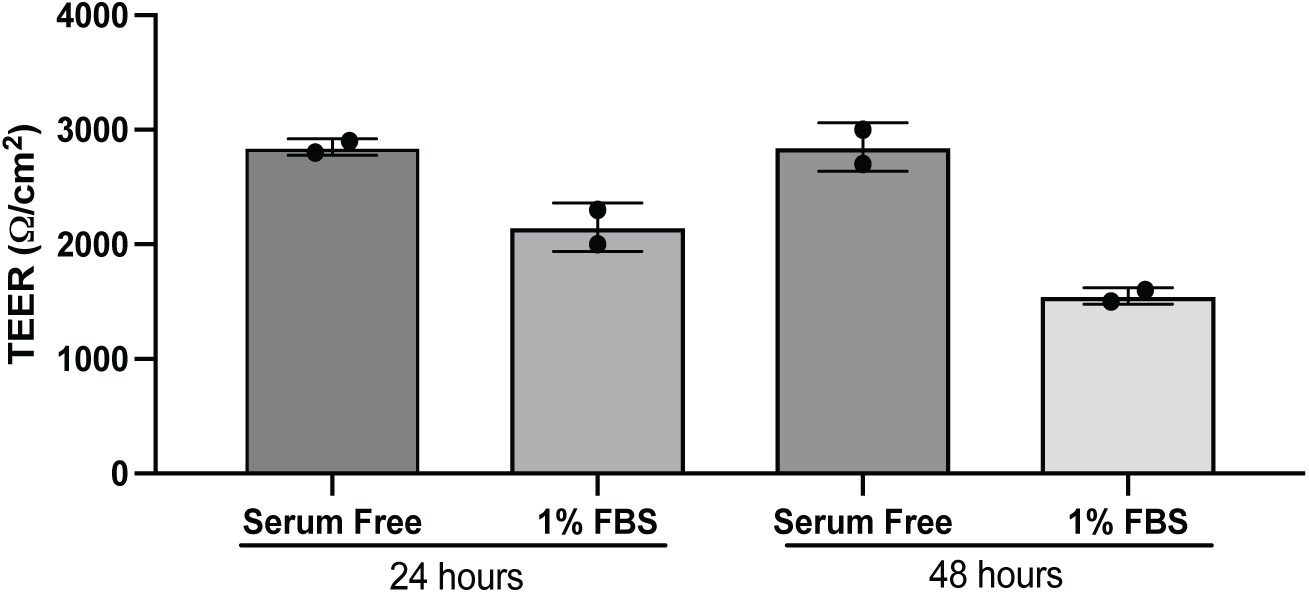
Fetal bovine serum reduces hPSC-derived BMEC barrier function. hPSC-derived BMECs were plated in Transwell plates and the media in the lower chamber was replaced with sfPCM with or without 1% FBS 48 hours after plating. TEER was measured at 24 or 48 hours after media replacement.

**Figure S4:**
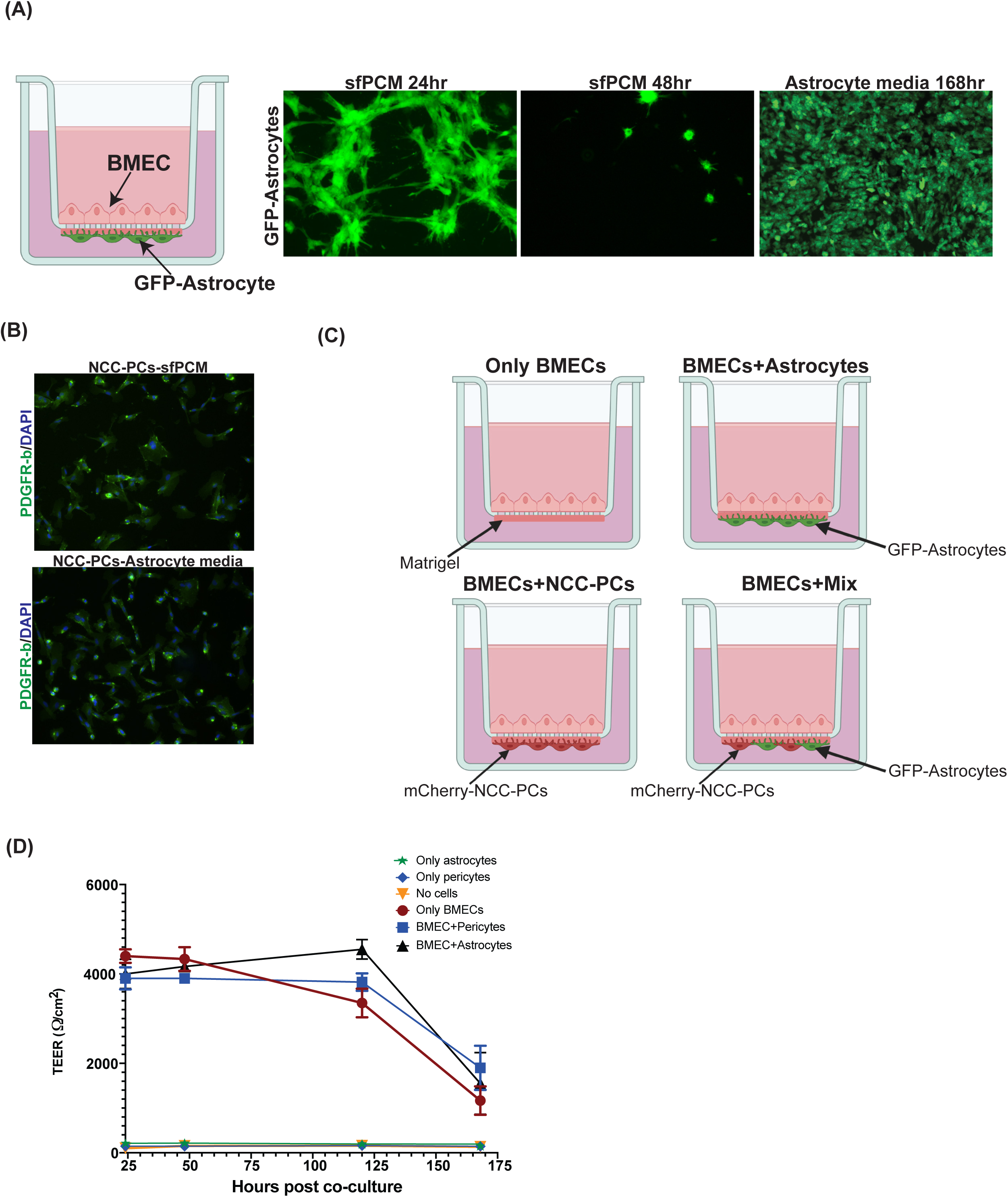
Optimization of the ccBBB model. **(A)** GFP-tagged astrocytes and BMECs were cultured in the ccBBB system with either astrocyte media of sfPCM in the lower chamber. At the indicated time after the initiation of co-culture GFP-astrocytes were imaged by florescence microscopy. **(B)** hPSC-derived NCC-PCs were cultured for 72 hours in either sfPCM or astrocyte media and then fixed and stained with an antibody against PDGF-R beta. **(C)** Schematic of conditions tested in the ccBBB system.

**Figure S5:**
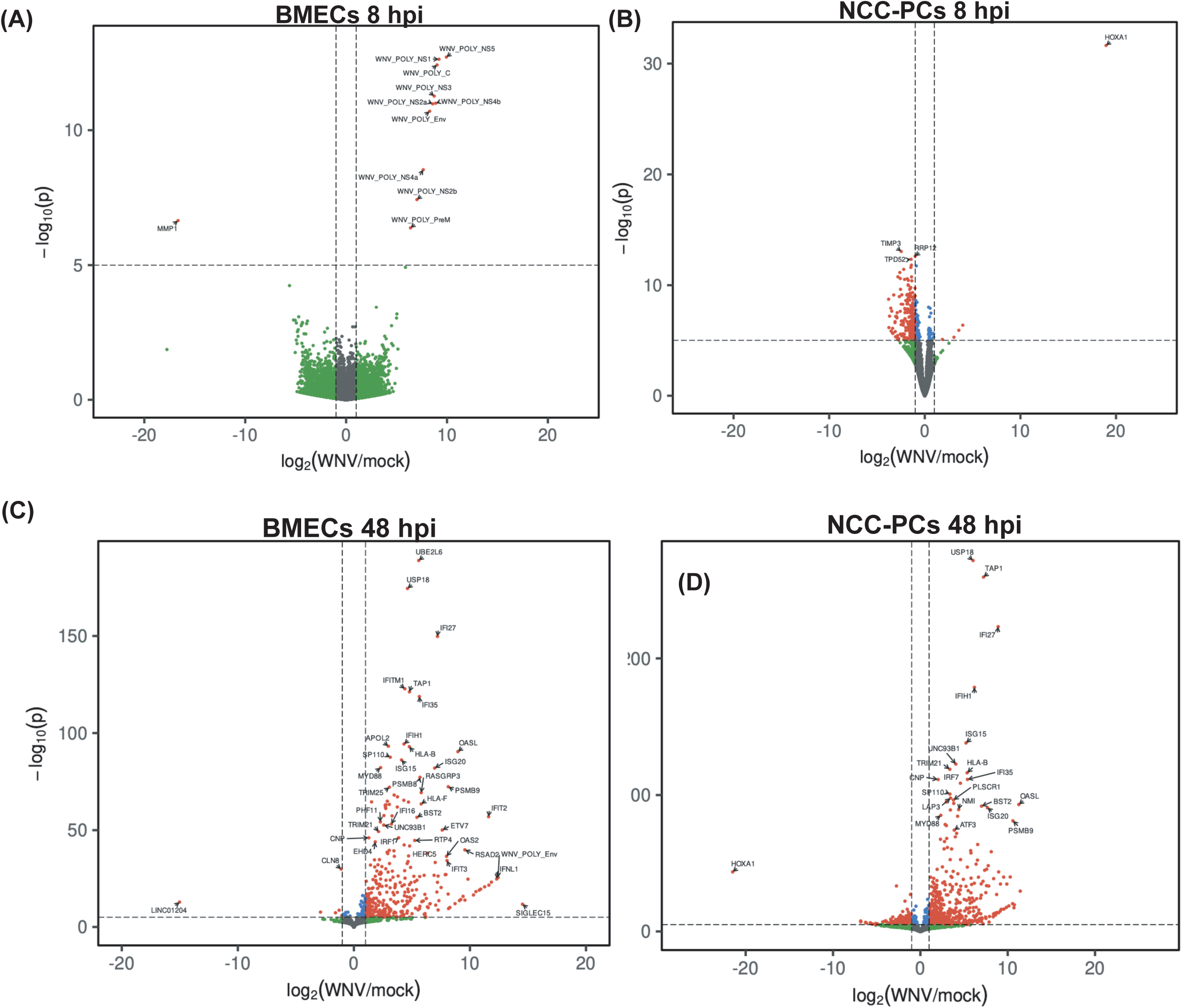
NCC-PCs and BMECs in the cBBB system display divergent transcriptional responses to WNV infection. hPSC-derived BMECs and NCC-PCs were plated in the ccBBB model as described in Fig. 3A. WNV was added to the apical chamber at an MOI of 1. Total cellular RNA was isolated from BMECs and NCC-PCs at the indicated time post infection and analyzed by bulk RNA sequencing **(A)** Volcano plot showing differential gene expression in mock versus WNV-infected BMECs at 8 hours post infection. **(B)** Volcano plot showing differential gene expression in mock versus WNV-infected NCC-PCs at 8 hours post infection. **(C)** Volcano plot showing differential gene expression in mock-versus WNV-infected BMECs at 48 hours post infection. **(D)** Volcano plot showing differential gene expression in mock-versus WNV-infected NCC-PCs at 48 hours post infection. All volcano plots are shown without empirical Bayes shrinkage applied to fold-change estimates^33^.

## Methods

### hPSC lines and maintenance

H1 (WA01) embryonic stem cells were obtained from WiCell. iPS11 induced pluripotent stem cells were obtained from Alstem. All hPSCs were maintained in feeder-free conditions in mTeSR Plus media (StemCell Technologies) on Matrigel (Corning) in 6-well tissue culture dishes. For passaging, hPSCs were detached as clumps using Versene Solution (Thermo Fisher Scientific) and replated at a ratio of 1:8-1:10.

### Directed differentiation of hPSCs to neural crest derived pericytes

H1 or iPS-11 hPSC were differentiated to Neural Crest cells using the Stemdiff Neural Crest Differentiation kit (Stem Cell Technologies Cat#08610) according to the manufactures protocol. At passage two neural crest cells were plated in Neural crest cell expansion media (NCC Expansion media, Table 2) on fibronectin coated plates. Neural crest cells were cryopreserved at passage three. For differentiation to neural crest derived pericytes neural crest cells were thawed directly into sfPCM (Table 2) on to fibronectin coated plates. Cells were cultured in sfPCM for a minimum of 10 days and passaged 1:3 when they reached 90% confluency. At day 10-14 cells were cryopreserved at 200,000 cells/ml.

**Table 2.**

|  | Final Concentration | Source | Catalog # |
| --- | --- | --- | --- |
| **MeIM** |  |  |  |
| E6 |  | Thermo Fisher Scientific | A1516401 |
| L-Ascorbic acid 2-phosphate sesquimagnesium salt hydrate (AA) | 60ug/ml | Sigma | A8960-5G |
| CHIR 99021 | 8uM | Biogems | 2520691 |
| BMP4 | 25ng/ml | Peprotech | 120-05ET |
| **NCC Expansion media** |  |  |  |
| DMEM/F12 |  | Thermo Fisher Scientific | 11320033 |
| B27 | 1:50 | Thermo Fisher Scientific | 17504044 |
| N2 | 1:100 | Thermo Fisher Scientific | 17502048 |
| L-glutamine | 1:100 | Thermo Fisher Scientific | A2916801 |
| NEAA | 1:100 | Thermo Fisher Scientific | 11140050 |
| EGF | 20ng/ml | Peprotech | AF-100-15 |
| FGF-2 | 20ng/ml | R&D Systems | BT-FGFB-GMP-025 |
| CHIR 99021 | 3uM | Biogems | 2520691 |
| **PC** |  |  |  |
| E6 |  | Thermo Fisher Scientific | A1516401 |
| Forskolin | 2nM | Biogems | 6652995 |
| AA | 60ug/ml | Sigma | A8960-5G |
| VEGF | 200ng/ml | Peprotech | 100-20-50µg |
| **PC2** |  |  |  |
| hESFM |  | Thermo Fisher Scientific | 11111044 |
| Forskolin | 2nM | Biogems | 6652995 |
| AA | 60ug/ml | Sigma | A8960-5G |
| VEGF | 200ng/ml | Peprotech | 100-20-50µg |
| **PC3** |  |  |  |
| hESFM |  | Thermo Fisher Scientific | 11111044 |
| AA | 60ug/ml | Sigma | A8960-5G |
| VEGF | 200ng/ml | Peprotech | 100-20-50µg |
| SB 431542 | 10uM | Biogems | 3014193 |
| **sfPCM** |  |  |  |
| hESFM |  | Thermo Fisher Scientific | 11111044 |
| B27 | 1:50 | Thermo Fisher Scientific | 17504044 |
| EGF | 20ng/ml | Peprotech | AF-100-15 |
| Heparin | 2ug/ml | StemCell Technologies | 07980 |
| SB 431542 | 10uM | Biogems | 3014193 |
| PDGFbb | 10ng/ml | Peprotech | 100-14B-10UG |

### Directed differentiation of hPSCs to mesoderm derived pericytes

All media recipes are described in Table 3. H1 hPSCs were cultured in E8 media (Thermo Fisher Scientific) and dissociated using Accutase (Thermo Fisher Scientific) into a single cell suspension and plated at 15,000 cells/cm^2^ in E8 media with Y-27632 (10μM). Y-27632 was removed after 24 hours and cells cultured in E8 media until they reached approximately 70% confluency at which point the media was replaced with MelM media (Table 2), this is considered day 0. On day 2 the media was replaced with PC/SMC1 media. On day 5 the media was replaced with PC/SMC2 media (Table 2) for 24 hours and then cells were passaged 1:3 on to fibronectin (FC01010MG, Fisher Scientific) coated tissue culture plates in PC3 media. Plates were coated with fibronectin at 10ug/ml for 30 minutes at 37°C prior to the addition of cells. After 48 hours media was replaced with sfPCM (Table 2) media. After 48 hours the media was replaced with fresh sfPCM (Table 2) and cells kept in culture for 7 days prior to freezing.

### Directed differentiation of hPSCs to brain microvascular endothelial cells (BMECs)

The differentiation of H1 hPSCs to brain microvascular endothelial cells was based on a previously published protocols^34–36^. Briefly, hPSCs were dissociated using Accutase (Thermo Fisher Scientific) into a single cell suspension and plated at 15,000 cells/cm^2^ in StemFlex media with Y-27632 (10μM). Y-27632 was removed after 24 hours and at 48 hours post-plating the media was replaced with Unconditioned Media (UM) (100ml Knock-out Serum Replacement (Thermo Fisher Scientific), 5ml non-essential ammino acids (Thermo Fisher Scientific), 2.5ml GlutaMax (Thermo Fisher Scientific), 5ml Pen/Strep (Thermo Fisher Scientific), 3.5μl β-mercapto-ethanol (Sigma), and 392.5ml DMEM/F12(1:1), this was considered day zero. The media was replaced daily with fresh UM. On day six the media was changed to hESFM (Thermo Fisher Scientific) supplemented with 20 ng/mL bFGF (Peprotech), 10 µM retinoic acid (RA) (Sigma), and 1:50 B27 (Thermo Fisher Scientific). The media was not changed for 48 hours. On day eight, cells were washed once with DPBS and incubated with Accutase for 30 minutes at 37°C. Cells were collected via centrifugation and plated onto either standard tissue culture plates or Transwell plates (Corning #3460). Both plates and Transwells were coated with 400 µg/mL collagen IV (Sigma Aldrich) and 100 µg/mL fibronectin (Fisher Scientific) overnight with collagen and fibronectin and washed 1X with PBS prior to the addition of cells. For tissue culture plates, cells were plated at a density of 250,000 cells/cm^2^, whereas for Transwell plates, cells were plated at a density of 1.1 x 10^6^ cells/cm^2^ on the Transwell membrane. bFGF and RA were removed from the medium 24 hours after plating.

### Virus propagation and titration

WNV (Nea Santa Greece 2010 Strain), JEV (India R53567), POWV (LB Strain) were obtained from BEI Resources. All three viruses were propagated in Vero E6 cells (ATCC CRL-1586) cultured in Dulbecco’s modified Eagle’s medium (DMEM) supplemented with 2% fetal calf serum (FCS), penicillin (50 U/mL), and streptomycin (50 mg/mL). Viral titers were determined in Vero E6 cells by plaque assay. Work with POWV and JEV was performed in the biosafety level 3 (BSL3) at the Ragon Institute (Cambridge, MA) following approved SOPs. For infection of BMECs, NCC-PCs, or Astrocytes the cell culture media was removed and replaced with inoculating virus diluted in a minimum volume of culture media. The cells were placed at 37°C for 1 hour at which point the inoculum was removed and replaced with sfPCM (NCC-PCs), hESFM+B72 (BMECs), or astrocyte media (astrocytes).

### Cell seeding in to ccBBB system

BMECs were plated in 12 well Transwells as described above. 72 hours after FGF/RA removal the inserts were removed from the 12 well plates and placed upside down onto the inverted lid of a Deep Well 6-well plates (Corning Cat#355467). Matrigel was added the lower side of the semipermeable membrane (opposite side as BMECs). The Transwell inserts were covered with the lid of the Deep Well 6-well plate to prevent evaporation and placed at 37C for 30 min. After incubation excess Matrigel was removed and 1.5e5 astrocytes or NCC-PCs resuspended in astrocyte media were added the Matrigel coated side of the Transwell insert. The Transwell inserts were again covered with the lid of the Deep Well 6-well plate and placed at 37C for three hours to allow cells to adhere. After incubation the Transwell inserts were placed back in 12-well plates and 0.5ml hESFM+B27 was added to the top chamber of the Transwell and 1.5ml of astrocyte media added to the lower chamber. The Transwell plate was kept at 37C until the time of infection.

### Flow cytometry

Cells were trypsinized with TrypLE Express for ∼5-7 minutes or until cells detached. Cells were washed once with PBS and resuspended in IC Fixation Buffer (Thermo Fisher) and incubated for 30 minutes at room temperature. Following incubation cells were washed twice in 1X Permeabilization Buffer (Thermo Fisher) and then resuspended in 100ul of 1X Permeabilization buffer containing the PDGFR-beta primary antibody at 1:100. Cells were incubated for 1hour at room temperature and then washed twice with 1X Permeabilization Buffer and then antibody incubation procedure was repeated using Alexaflour 488 secondary antibody at 1:200. After antibody incubation cells were washed twice in Permeabilization Buffer and resuspended in 1ml of Flow Cytometry Buffer (PBS + 1% BSA) Stained cells were analyzed on a BD Fortessa UV/Blue/Green/Red 4-laser flow cytometer. Cells were isolated as a population and then selected for singlets before setting gates to determine positive populations.

### Immunofluorescence microscopy

Cells were fixed with 4% paraformaldehyde for 15 minutes at room temperature. Fixed cells were permeabilized for 15 minutes using 0.1% TX-100. Cells were then blocked with 3% BSA in PBS for 1 hour at room temperature. Cells were incubated with primary antibodies (see Table 2) diluted in antibody buffer (0.01%TX-100 and 1%BSA in PBS) overnight at 4°C. Cells were washed with 0.05% Tween in PBS and incubated for 1 hour at room temperature with secondary antibodies (see Table 2) diluted in antibody buffer. FITC-Phalloidin was added at 1:2000 and DAPI at 1:5000 with secondary antibodies.

### Measurement of trans-endothelial electrical resistance

BMECs were plated onto three Transwell filters (Corning #3460) using the procedure described in “Directed Differentiation of hPSCs to brain microvascular endothelial cells” ^34^ .48 hours after plating the media was removed from the apical side of the Transwell and replaced with SMC-conditioned media. At each time point, the TEER was measured in three separate wells. All data are represented as mean ± standard deviation for these collective measurements. Following the medium change on day 0 of subculture to remove bFGF and RA, no further medium changes were performed for the duration of the experiments.

### FITC-Dextran Permeability Assay

Brain microvascular endothelial cells were seeded in 12-well Transwell plates (Corning #3460) at 1.1e6 cells/transwell on day 8 of the differentiation protocol . bFGF and RA were removed from the medium 24 hours after plating. 48 hours after plating Transwells were placed in the ccBBB system (see above) with either hPSC-derived astrocytes, hPSC-derived NCC-PCs, or primary pericytes. xx hours after the initiation of co-culture the media on the basolateral side was replaced with 1ml of fresh BMEC media. FITC-tagged dextran (10 μM) was suspended in 0.5 ml of BMEC medium and added to the apical side of the Transwell. To determine the level of transcytosis following 2 hours of incubation in a 37°C incubator on a rotating platform, we collected 150 μl from the 1ml of EC medium on the basolateral side of the Transwell. Fluorescence intensity in the collected media was measured using a SpectraMax iD3 plate reader set to 495-nm excitation and 519-nm emission settings.

### Bulk RNA-sequencing

RNA was extracted from cells using the RNeasy Plus Mini kit (QIAGEN) following the manufacturer’s protocol. For cell characterization samples (Figure 1, Supplemental Fig S1-S2) libraries were prepared using the Swift RNA library prep kit (Swift Biosciences).

For ccBBB sample sequencing (Figure 4, Supplemental Fig. S5) libraries were prepared using the SMART-Seq® v4 Ultra® Low Input RNA Kit. All samples were sequenced on a NovaSeq 6000 sequencer.

### Analysis of gene expression

Paired-end reads (51x 51 bp) were mapped to a reference genome using the STAR aligner (v. 2.7.1a)^37^. The reference genome was a composite of the human genome (GRCh38) and that of the West Nile virus (Nea Santa Greece 2010, GenBank HQ537483.1). Counts for human protein coding genes and lncRNAs (ENSEMBL release 93 annotations), together with the viral genes, were tabulated using featureCounts^38^. Differential expression of mRNAs was assessed for pairwise contrasts between conditions using estimated fold-changes and the Wald statistic in DESeq2 (v. 1.36.0)^32^. From exploratory analysis, within-condition variance was observed to be relatively uniform across conditions, so per-gene dispersions were estimated using all conditions. Unless otherwise noted, empirical Bayes shrinkage ^33^ was applied to fold-change estimates. Gene-set enrichment analysis was carried out between pairs of conditions using GSEA ^26^(v. 4.1.0) with counts that were normalized by DESeq2 and 1000 gene-set permutations were used to estimate p-values. Gene-sets that were tested for enrichment came from the Hallmark collection^39^ at the MSigDB.

### Cell type characterization from bulk RNA sequencing and PCA

RNA sequencing reads were mapped to the human genome (GRCh38) using the STAR aligner (version 2.7.1a). The mapped reads were assigned to the human genes (ENSEMBL release 101 annotations) using featureCounts with the parameters ‘-p -s 2’, suitable for paired-end reversely-stranded libraries. Before using the ‘prcomp’ function in R (version 4.2.1) to perform PCA, the normTransform function of DESeq2 (version 1.36.0)^32^ was used to normalize, center and log-transform (with a pseudocount of 1) the counts matrix.

